# Menstrual cycle and hormonal contraceptive intake alter the electrophysiological signature of selective auditory spatial attention

**DOI:** 10.64898/2026.07.29.741520

**Authors:** Michael-Christian Schlüter, Marlies Pinnow, Martin Brüne, Jörg Lewald

**Affiliations:** Auditory Cognitive Neuroscience, Department of Cognitive Psychology, Faculty of Psychology, Ruhr University Bochum, Bochum, Germany; Motivation Lab, Department of Biopsychology, Faculty of Psychology, Ruhr University Bochum, Bochum, Germany; LWL University Hospital, Department of Psychiatry, Psychotherapy and Preventive Medicine, Division of Social Neuropsychiatry and Evolutionary Medicine, Ruhr University Bochum, Bochum, Germany

**Keywords:** menstrual cycle, sex differences, selective auditory spatial attention, cocktail-party listening, sound localization, auditory evoked potentials

## Abstract

The fluctuations in steroid hormones associated with the female menstrual cycle have been suggested to modulate attentional brain processes. This study focused on cycle-dependent changes in the electrophysiological correlates of sound localization in a cocktail-party task, which has been shown to reflect the neural correlates of auditory selective attention. Event-related-potentials were recorded in three (1 - luteal, 2 - menses, 3 - follicular) phases (determined based on self-reports) in women not taking hormonal contraceptives, at three corresponding points in time in women taking oral hormonal contraceptives, and in men. An effect of phase in women was obtained exclusively for the N2 component, which may reflect processes of auditory selective spatial attention. N2 amplitudes were strongest in phase 1 (luteal phase or initial phase of pill intake) and lowest in phase 2 (menses or pause of pill intake). Compared with the male group, the N2 amplitudes were lower in women in phase 2, but not in phase 1 and 3. Cortical source localization at the time of the N2 revealed a related modulation of electrical activity in right superior frontal sulcus depending on phase. Also, electrical activity in left inferior frontal gyrus was increased in women in phase 2 compared with men. These findings suggest that the neural correlates of auditory selective spatial attention varied in women depending on their hormonal phase. The observed pattern is compatible with the view that higher concentrations of female steroid hormones enhance brain processes of selective spatial attention.

## 1 Introduction

Localizing and identifying a speaker of interest among multiple interfering sound sources in a cocktail party situation (Cherry, 1953) is often a demanding challenge in normal life. Nevertheless, the brain mechanisms of auditory selective attention are usually capable of automatically directing the focus of interest to the relevant sound source without conscious effort, while irrelevant sounds are suppressed through object-based auditory attention mechanisms (Alain & Arnott, 2000; Shinn-Cunningham, 2008; Zion Golumbic et al., 2013). Several studies on this topic have reported a male advantage in sound localization in an auditory scene with multiple sound sources, thus suggesting a sex difference for brain processes of auditory selective spatial attention (Zündorf et al., 2011; Lewald & Hausmann, 2013; Plass, 2013). Similarly, advantages for men were also found for other audiospatial tasks, such as monaural vertical sound localization (Lewald, 2004), perception of sound motion (Schiff & Oldak, 1990; Neuhoff et al., 2009; Grassi, 2010), detection of spatial oddballs (Simon-Dack et al., 2009), discrimination of interaural level differences (Wright & Dai, 2022), and perception of binaural beats (Tobias, 1965). In contrast, female advantages have been reported for non-spatial auditory performance, such as hearing sensitivity, signal detection in the presence of a masker, detection of a temporal gap in noise (McFadden et al., 2006; Gold et al., 2015), and speech recognition (e.g., Lutman, 1991; Wiley et al., 1998; for review, see McFadden, 1998; Halpern, 2012). Sex differences were found also using electrophysiological and neuroimaging methods. For example, auditory brainstem responses were generally stronger in women than in men (Jerger & Hall, 1980; for review, see Aloufi et al., 2023). Also, stronger P300 components of the auditory event-related potential (ERP) have been shown in women (for review, see Melynyte et al., 2018; Bianco et al., 2022). Furthermore, functional magnetic resonance imaging (fMRI) with audiospatial tasks (e.g., high-versus low-confidence spatial memory) indicated stronger activation in men than in women in various cortical areas, in particular in lateral prefrontal cortex (Alain et al., 2001; Spets & Slotnick, 2020; Spets et al., 2021; Slotnick, 2021; Sneider & Silveri, 2021). It has been suggested that some of these auditory sex differences may be related to findings in the visual domain, thus reflecting multi-or supra-modal, rather than unisensory, phenomena. The above-mentioned male advantage in cocktail-party sound localization (Zündorf et al., 2011; Lewald & Hausmann, 2013; Plass, 2013; Lewald, 2026) has been associated with studies on sex differences in visuospatial cognition, in particular the well-documented male advantage in the Mental Rotation Test (MRT; Vandenberg et al., 1978; Voyer et al., 1995; Kimura, 1996; Postma et al., 1999; Hausmann et al., 2000; Maki et al., 2002; Hirnstein & Hausmann, 2009; Lippa et al., 2010; Maeda et al., 2013; Kasai 2021).

At this point, it is important to note that more recent studies in the visuospatial domain provided evidence that sex differences depend on the female menstrual cycle, thus suggesting modulation of the underlying brain processes by sex hormone concentrations (e.g., Hampson, 1990a, 1990b; Silverman et al., 1993, 1999; Hausmann et al., 2000; McCormick & Teillon, 2001; Schöning et al., 2007; Simić & Santini, 2012; Scheuringer et al., 2017). Hausmann et al. (2000) obtained high MRT scores in women during their menstrual phase and low scores during their midluteal phase. On the basis of analyses of hormone concentrations, these authors concluded that estradiol had a negative influence on MRT performance and testosterone a positive one. Accordingly, McCormick and Teillon (2001) reported poorer MRT performance of women in the luteal phase, that is, when levels of progesterone and estradiol are high. Also, Simić et al., (2001) and Santini (2012) found best MRT performance during the phases of menstrual cycle in which levels of sex hormones were low. In the fMRI study by Schöning et al. (2007), brain activation with mental rotation in women was correlated with estradiol in early follicular and midluteal cycle phases in frontal and parietal cortical areas and with testosterone in the early follicular phase (Gizewski et al., 2006; Halari et al., 2006). In men, MRT performance was better with high testosterone levels (Hooven et al., 2004). Also, a correlation was identified between follicle-stimulating hormone (FSH) levels and men’s performance in spatial tasks: higher FSH levels led to poorer performance, whereas lower FSH levels enhanced performance (Gordon & Lee, 1986). Thus, levels of female and male sex hormones may influence visuo-spatial performance in different ways.

These relations between measures of visuo-spatial cognition and hormonal phase raised the question of whether similar effects may also exist in the auditory domain. So far, however, almost nothing is known about this topic. Only Walpurger et al. (2004) investigated effects of the menstrual cycle on the auditory ERP. In a non-spatial oddball task, N2 latency was found to be shorter during menses compared to the follicular and luteal phases, suggesting that low levels of female sex hormones may strengthen attentional processing in cortex (Walpurger et al., 2004). Here, we addressed this problem using a spatial cocktail-party task, as used in previous research (e.g., Lewald & Getzmann, 2015; Lewald et al., 2016; Hanenberg et al., 2019, 2021), which required the listener to identify a relevant sound source by analyzing spatial information while simultaneously filtering out distracting sources in a complex acoustic scene. Using cocktail-party paradigms, the N2 component of the ERP has been identified as a specific correlate of selective spatial attention (e. g., Hill & Miller, 2010; Gamble & Luck, 2011; Lewald & Getzmann, 2015; Lewald et al., 2016; Hanenberg et al., 2019, 2021). Specifically, larger N2 amplitudes have been linked to enhanced neural processes underlying auditory attention. (Hanenberg et al., 2019, 2021). Thus, this approach seemed well-suited for investigating menstrual-cycle effects on attentional brain processes. Based on the previous studies mentioned above, the N2 was defined as the main electrophysiological measure. We hypothesized that the N2 amplitude will be largest during high-hormone phases and smallest during the low-hormone phases. Other ERP components as well as N2 latency were analyzed as secondary measures to determine whether phase-related modulations were specific to the N2 amplitude or reflected a more general change across the auditory ERP waveform.

For this purpose, normally cycling women not taking hormonal contraceptives (NHC) were recruited as participants. In addition, women taking oral hormonal contraceptives (HC) were included to also examine effects of regular intake and discontinuation of synthetic steroids (D’Souza et al., 2023). Finally, a group of men served as a reference. NHC women were tested at three points of their cycle (luteal phase; menses; follicular phase), HC women at three related points during regular intake of pills and discontinuation of intake (cf. Mihm et al., 2011; Reed & Carr, 2018; for details, see 2.4) and men once. The measurement points in time were chosen such that women were tested during maximum progesterone and intermediate estradiol concentrations (phase 1), low levels of both hormones (phase 2), and maximum estradiol and low progesterone levels (phase 3). The phases were determined based on the participants’ self-reported menstrual cycle information, without hormone measurements, since this approach is an established method for identifying phases with high and low hormone concentrations with sufficient accuracy (Sundström-Poromaa & Comasco, 2014; Allen et al., 2016).

## 2 Methods

### 2.1 Participants

Data were collected from a total of 100 young adult participants subdivided into 3 groups: *(1)* men (*n* = 35; mean age 25.20, SE 0.65, range 19-33 years); *(2)* normally cycling women not taking hormonal contraceptives (NHC; *n* = 27; mean age 22.93, SE 0.76, range 18-32 years); and *(3)* women taking oral hormonal contraceptives (HC; *n* = 38; mean age 22.00, SE 0.48, range 18-30 years). All contraceptive pills used were combined estrogen-progesterone oral contraceptives, with typical estrogen and progesterone doses of about 20-30 µg/d. Menstrual cycle phase was classified using participants’ self-reported menstrual history, cycle regularity, and the timing of testing relative to the onset of menstruation (for details, see 2.4).

All participants were right-handed (mean laterality quotient 90.74; SE 1.47, range 30-100), as determined by the 10-item version of the Edinburgh Inventory (Oldfield, 1971). Hearing thresholds of all participants were within the normal range, as assessed by standard pure-tone audiometry in the frequency range from 125 Hz to 8 kHz (mean hearing level ≤ 25 dB; Oscilla USB100, Inmedico, Lystrup, Denmark). Further inclusion criteria were no pregnancy, confirmed by a negative pregnancy test at each session, and self-reported constant menstrual cycle for female participants, as well as no intake of neuroactive substances, no neurological or psychiatric illnesses, and an age between 18 and 35 years for all participants. Data of 20 further participants were discarded. These participants were either dropouts (*n* = 9) or did not meet the inclusion criterion of at least 40% correct responses (*n* = 4), or the quality of EEG recording was insufficient (*n* = 7).

As no directly comparable studies were available, a target sample of 30 completers per group (plus anticipated dropouts) was chosen based on previous studies reporting effects of sex on auditory spatial scene analysis (*n* ≤ 27; Zündorf et al., 2011; Lewald & Hausmann, 2013) and effects of the menstrual cycle on auditory functions (Kirschbaum et al., 1999; Walpurger et al., 2004; Zhu et al., 2016; Souza et al., 2017; Emami et al., 2018; Morris et al., 2019; McFadden et al., 2021). In addition, a sample-size estimation was conducted using G*Power Version 3.1.9.2 (Faul et al., 2007), assuming effect sizes (Cohen’s *d*) between 0.6 and 0.7 (α = 0.05, 1–β = 0.80) for the main effect of phase on the N2 amplitude in the repeated-measures ANOVA (see 3.2.1), based on effect sizes for sex differences in selective attention reported by Zündorf et al. (2011) and Lewald and Hausmann (2013).

This calculation resulted in estimated group sizes of 26–36 participants. Because a larger number of dropouts was anticipated, we aimed to recruit a total sample of 120 participants (40 participants per group). The sample-size estimation was based on previously reported between-group sex effects and was used to inform the recruitment target. It was therefore not designed to establish 80% power for the within-participant phase effect examined in the primary repeated-measures analysis, and the resulting sample size should not be interpreted as demonstrating adequate power for all repeated-measures or mixed-model effects investigated in the present study.

This study conformed to the Code of Ethics of the World Medical Association (Declaration of Helsinki) published in the British Medical Journal (July 18, 1964) and was approved by the Ethics Committee of the Medical Faculty of the Ruhr University Bochum. All participants gave their written informed consent to take part in the study. They received either course credits or were paid for participation.

### 2.2 Apparatus

The apparatus and the general procedure used here were similar to those described for the multiple-sources condition in Lewald and Getzmann (2015). Experiments were conducted in a dimly illuminated, anechoic, and sound-proof room (for details, see Guski, 1990). The participant sat in a vertically adjustable chair. The position of the participant’s head was held constant by a chin rest. In front of the subject, a semicircular array of 91 broad-band loudspeakers (SC 5.9, Visaton, Haan, Germany) was mounted in the horizontal plane at a distance of 1.5 m from the centre of the participant’s head at ear level. The loudspeakers covered the frontal 180° range from-90° (left) to 90° (right) in steps of 2° with the centre loudspeaker at 0° (for a more detailed description, see Lewald et al., 2004). In the present study, only four loudspeakers located at-90°,-30°, 30°, and 90° were used. A red light-emitting diode (LED; diameter 3 mm, luminance 0.025 mcd) located immediately below the central loudspeaker served as fixation target and was always on.

### 2.3 Auditory stimuli and task

Four different animal vocalizations (‘birds chirping’; ‘dog barking’; ‘frog’; ‘sheep’), taken from an online sound library (Marcell et al., 2000) and set to a constant duration of 600 ms (identical onset, without digital manipulation of duration) and identical peak levels using Cool Edit 2000 software (Syntrillium Software Corporation, Phoenix, AZ, USA), were used as auditory stimuli. Details of these stimuli have already been described in Lewald and Getzmann (2015) and Lewald (2016). For each animal, four different versions of the vocalizations were generated by choosing different cut-outs from the original recording. Sound files were digitized at a sampling rate of 48 kHz and a resolution of 16 bits, and converted to analog form via a PC-controlled sound card (Terrasoniq TS88 PCI, TerraTec Electronic, Nettetal, Germany). In each trial, vocalizations of the four different animals were presented simultaneously, each emitted from a different loudspeaker with simultaneous onset. The combination of the four animal voices had an overall level of 66 dB(A).

The vocalization ‘frog’ was always defined as the target stimulus, since it was most easily recognizable compared with the other three animal voices, which were used as distractors. Prior to the experiment, the participants were informed that there were four possible sound locations: far to the left, front left, front right, and far to the right. The task was to localize the target sound by pressing one out of four response buttons (corresponding to the four possible target locations) on a response box immediately after each stimulus presentation (for further details, see Lewald & Getzmann, 2015). The response box was held in the left hand, and the participants responded with their right index finger. To minimize EEG alpha-activity and eye-movements, participants were instructed to focus on the central fixation LED. Prior to the experiment, participants were familiarized with the task in a few practice trials. Locations of target and distractor stimuli changed between trials in a fixed random order. Stimuli were presented with an interstimulus-interval of 2.4 s (3 s trial duration). Participants had to respond within 1.9 s after stimulus offset. The experiment comprised 576 trials, with one-time presentation of each possible combination of target/distractor locations and versions of animal vocalizations. After 288 trials, participants were allowed to rest for a few minutes. The timing of the stimuli and the recording of the participants’ responses were controlled by custom-written software. No feedback was given to the participants at any time.

### 2.4 Study design

NHC participants were tested in three sessions timed to coincide with the following cycle phases (determined by self-report): luteal phase (phase 1; 7-13 days before the anticipated onset of the menstrual cycle); menses (phase 2; 1st to 5th day of bleeding); and follicular phase (phase 3; 7-13 days after the onset of the menstrual cycle). For phase 1, the day of onset of the subsequent menstrual period was used retrospectively. As argued by Walpurger et al. (2004), the known variability of cycle duration is mainly confined to the follicular phase with the onset of the new menses beginning about 14 days after ovulation (Lein, 1979), such that the intervals for different phases can be counted backwards from the first day of the expected onset of the next menses.

HC participants were also tested in three phases: 7-13 days before the anticipated onset of the withdrawal bleeding (phase 1); on day 1 to 5 of withdrawal bleeding (phase 2); 7-13 days after the onset of the withdrawal bleeding (phase 3). Because of the chemical composition of the contraceptives used (synthetic ethinyl estradiol and synthetic progesterone/progestin), the blood concentrations of estradiol and progesterone are known to be continuously maximal during phases 1 and 3, while only low levels of these two hormones are present around the time of withdrawal bleeding, i. e., in phase 2 (e.g., Ekenros et al., 2022; Hirschberg, 2022).

Because of the hormonal composition of the contraceptives used (synthetic ethinyl estradiol and progestin), in the HC group, phases 1 and 3 corresponded to periods of active hormonal intake, whereas phase 2 corresponded to the hormone-withdrawal interval associated with withdrawal bleeding. For the HC group, the three selected phases thus represented periods during which exposure to synthetic sex steroids was relatively higher or lower.

In NHC and HC groups, the sequence of the three phases was counterbalanced across participants. Men were tested in only one session, since we assumed that potential effects of multiple testing might be negligible and comparisons with female groups in each of the three phases are justified.

### 2.5 ERP recording and analysis

ERP recording was identical to the methods described in Hanenberg et al. (2021). The continuous EEG was sampled at 1 kHz using a QuickAmp-72 amplifier (Brain Products, Gilching, Germany) and 58 Ag/AgCl electrodes, with positions based on the International 10-10 system. Four additional electrodes around the left and right eyes were used to record horizontal and vertical electro-oculograms. The ground electrode was positioned on the center of the forehead, just above the nasion, and two further electrodes on the left and right mastoids. Electrode impedance was kept below 5 kΩ. Raw data were band-pass filtered off-line (cut-off frequencies 0.5 and 25 Hz; slope 48 dB/octave), re-referenced to the average of 58 channels (56 EEG and 2 mastoid electrodes), and segmented into 2000-ms stimulus-locked epochs covering the period from 200 to 1800 ms relative to the onset of the sound stimulus.

The procedure of Gratton et al. (1983) was used for ocular artefact correction. Individual epochs exceeding a peak-to-peak amplitude of 200 µV were excluded from further analysis, using the automatic artefact rejection implemented in the BrainVision Analyzer software (Version 2.0; Brain Products, Gilching, Germany). The remaining epochs were baseline-corrected to a 200ms pre-stimulus window and averaged for each participant, separately for each session.

As in earlier studies using the same paradigm, only trials with correct responses were included in ERP analyses (e.g., Lewald & Getzmann, 2015; Lewald et al., 2016; Hanenberg et al., 2019, 2021). Thus, the results may reflect electrophysiological correlates of successful localization of the target sound, while minimizing potential effects arising from differences in error rates (Lewald & Getzmann, 2015).

The N2 was chosen a priori as the primary outcome variable since the central theoretical question concerned selective spatial auditory attention. Consequently, the primary confirmatory analysis was the effect of menstrual cycle phase on N2 amplitude. The peak of N2 was defined as the maximum negativity within 240–340 ms at FCz after sound onset. To assess whether phase-related effects were present at earlier or later stages of processing outside the temporal window of the N2, the P1 (10–110 ms after sound onset at Cz), N1 (60– 160 ms after sound onset at Cz), P2 (155–255 ms after sound onset at Cz) and LPC (400-700 ms after sound onset at Pz) components of the ERP were included in the analyses as secondary outcomes. Statistical inference for these outcomes was treated with due care and was separated from the primary N2 analysis. Similarly, comparisons between phases and comparisons between sexes were also conducted in secondary analyses. For each of the five components, peak-amplitude and latency data were submitted to statistical analyses. First, a two-factor (3 × 2) ANOVA with the within-participant factor phase (1; 2; 3) and the between-participants factor group (NHC; HC) was performed, in order to detect potential differences between groups and effects of phase. If this analysis indicated any group differences, each group was compared with data obtained in men using *t*-tests. If there were no differences between NHC and HC groups, data from both groups were collapsed and women were treated as one group for comparison with men. Comparisons between as well as pooling of the two female groups may be justified, as Petersen et al. (2014) found only marginal differences in estrogen and progesterone levels in NHC and HC women at three related phases.

### 2.5 Cortical source localization

The cortical sources of ERPs were localized using standardized low-resolution brain electromagnetic tomography (sLORETA; Pascual-Marqui, 2002), which is part of the LORETA-KEY software package (v20171101) of the KEY Institute for Brain-Mind Research, Zurich, Switzerland. Data were baseline corrected to a 200-ms pre-stimulus window for each participant, separately for each group. We employed sLORETA within 5-ms time windows around the individual ERP peak-amplitude values of N2 for each participant, with the individual latencies taken from the ERP analyses described above.

## 3 Results

### 3.1 Behavioral Results

The mean percentage of correct responses was clearly above chance level (25%) for each group (NHC: mean 90.15%, SE 2.24, range 46.76-99.77, *t*[26] = 29.12, *p* < 0.0001, *d* = 5.60; HC: mean 89.90%, SE 2.07, range 44.97-99.59, *t*[37] = 31.30, *p* < 0.0001, *d* = 5.08; men: mean 91.24%, SE 2.57, range 40.79-99.82, *t*[34] = 25.74, *p* < 0.0001, *d* = 4.35; Fig. 1). For NHC and HC groups, a two-factor (3 × 2) ANOVA including the within-participant factor phase (1; 2; 3) and the between-participants factor group (NHC; HC) did not reveal any significant main effect or interaction (*F* ≤ 0.96, *p* ≥ 0.39). Since there was no indication of a difference between NHC and HC groups, results obtained in men were compared with the group of women (NHC and HC collapsed) using *t*-tests. These *t*-tests indicated no significant sex differences at any phase (*t* ≤ 1.04, *p* ≥ 0.30). Thus, taken together, there was neither a modulation of performance as a function of time in NHC and HC groups, nor was there any difference between NHC and HC groups or between women and men. As already mentioned above (cf. 2.4), it has to be emphasized in this context that the paradigm used was optimized for detecting differences in ERP responses when the target was correctly localized, and that the analysis of error rates was not the primary focus of this study.

**Fig. 1.**
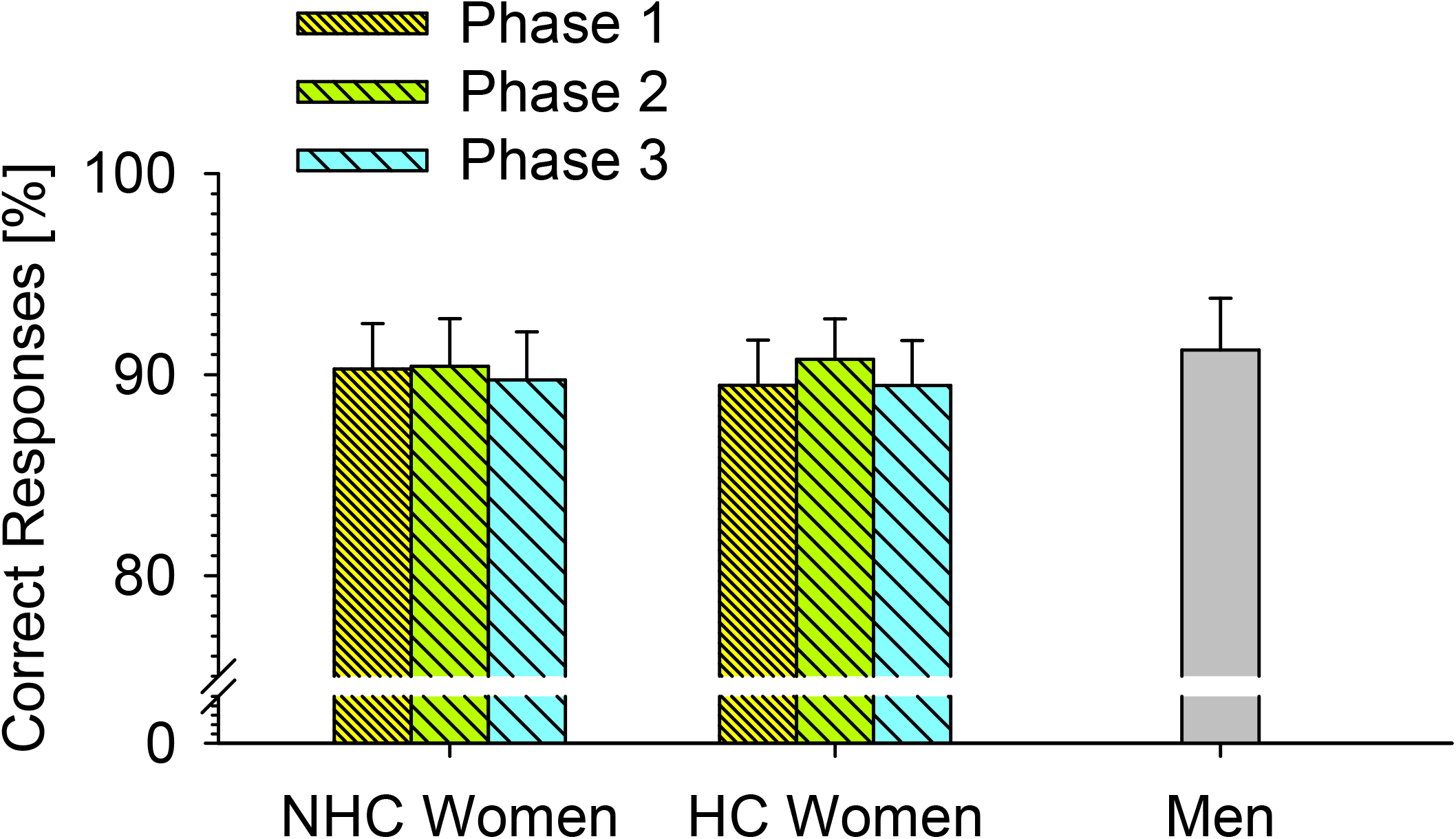
Behavioral results. Rates of correct responses are shown for NHC and HC women at three corresponding phases and for men. Error bars indicate standard errors across participants.

### 3.2 ERP components

In all groups, sound onset elicited a prominent response at vertex position Cz, (Fig. 2) mainly consisting of a positive deflection (P1), a negative deflection (N1), a second positive deflection (P2), a second negative deflection (N2), and a third positive deflection (LPC). The P2 and N2 waves were more positive (larger P2 and lower N2) in women than in men, with maximum sex differences at phase 2. Averaged across groups and conditions, mean latencies (with reference to sound onset) were 63.1 ms for P1 (SE 1.3 ms), 116.9 ms (SE 1.2ms) for N1, 201.9 ms (SE 1.7 ms) for P2, 286.6 ms (SE 2.0 ms) for N2 waves, and 461.0 ms (SE 7.3 ms) for the LPC.

**Fig. 2.**
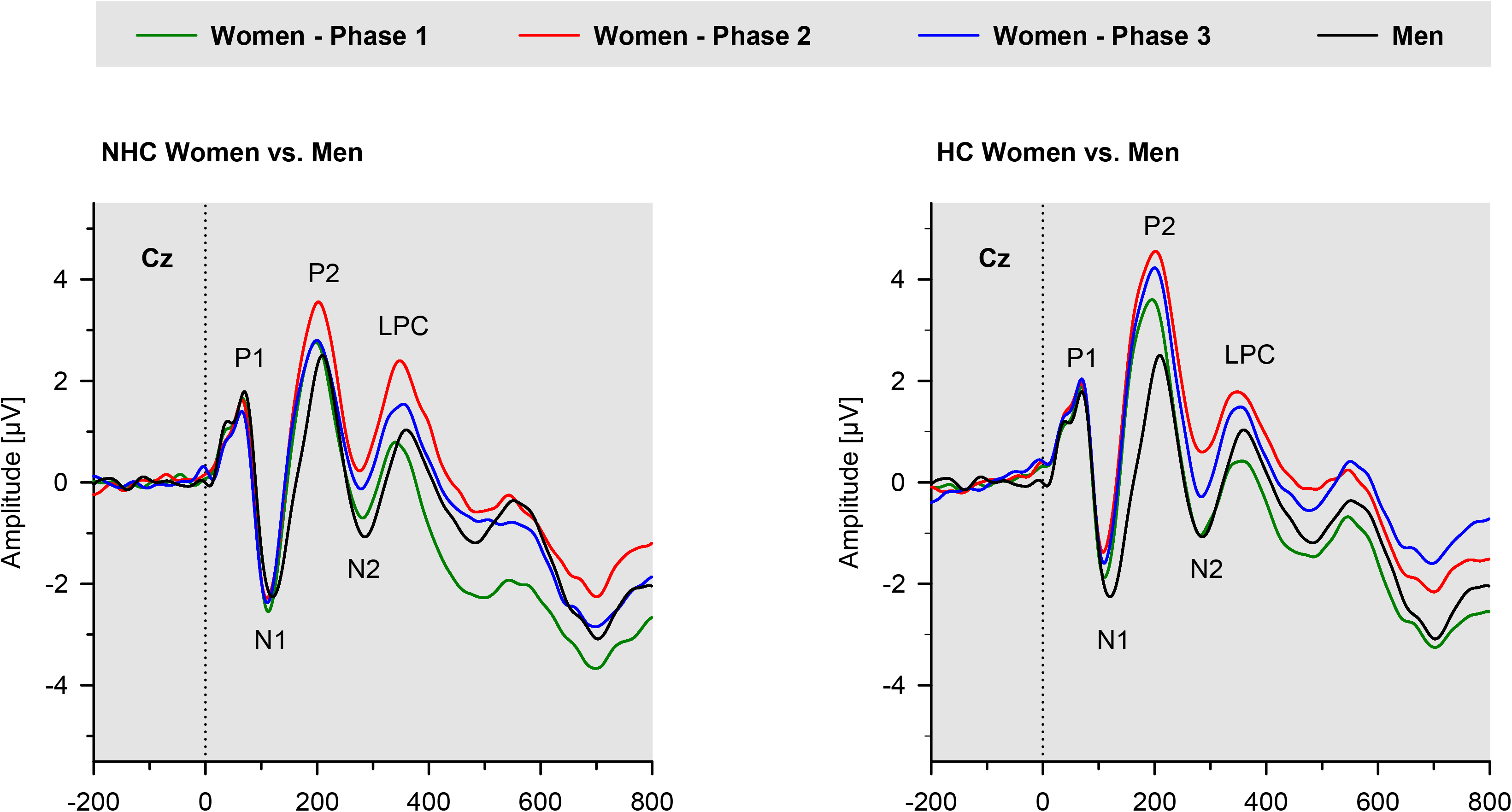
Grand-average ERPs for NHC (left panel) and HC women (right panel) compared with men. Waveforms at the Cz electrode are plotted as a function of time relative to the stimulus onset for women in phases 1, 2, and 3 and for men (tested once). ERP components P1, N1, P2, N2, and LPC are indicated in each panel.

For peak amplitudes and latencies of each of the five ERP components analyzed (P1; N1; P2; N2; LPC), a two-factor (3 × 2) ANOVA with the within-participant factor phase (1; 2; 3) and the between-participants factor group (NHC; HC) was first performed, in order to detect potential differences between groups and effects of phase. If this analysis indicated any group differences, each group was compared with data obtained in men using *t*-tests. If there were no differences between NHC and HC groups, data from both groups were collapsed and women were treated as one group for comparison with men.

#### 3.2.1 N2 wave

For the N2 peak amplitude at electrode position FCz, which was analyzed as the primary outcome measure, the two-factor ANOVA for NHC and HC groups at three phases showed a significant main effect of phase (*F*[2,126] = 3.43, *p* = 0.035, *h*_p_^2^ = 0.052), but no further significant results (*F* ≤ 0.50, *p* ≥ 0.48; Fig. 3). Post-hoc testing using paired *t*-tests revealed significantly stronger N2 amplitude at phase 1 compared with phase 2 (*t*[64] = 2.89, *p* = 0.005, *d* = 0.36), but no significant results for the two other comparisons (|*t*| ≤ 1.79, *p* ≥ 0.08; Bonferroni-corrected α = 0.017). Furthermore, the comparisons of men and women showed significantly larger N2 amplitudes in men than in women in phase 2 (*t*[94.83] = 2.49, *p* = 0.015, *d* = 0.46), but not in the other phases (*t* ≤ 1.60, *p* ≥ 0.11).

**Fig. 3.**
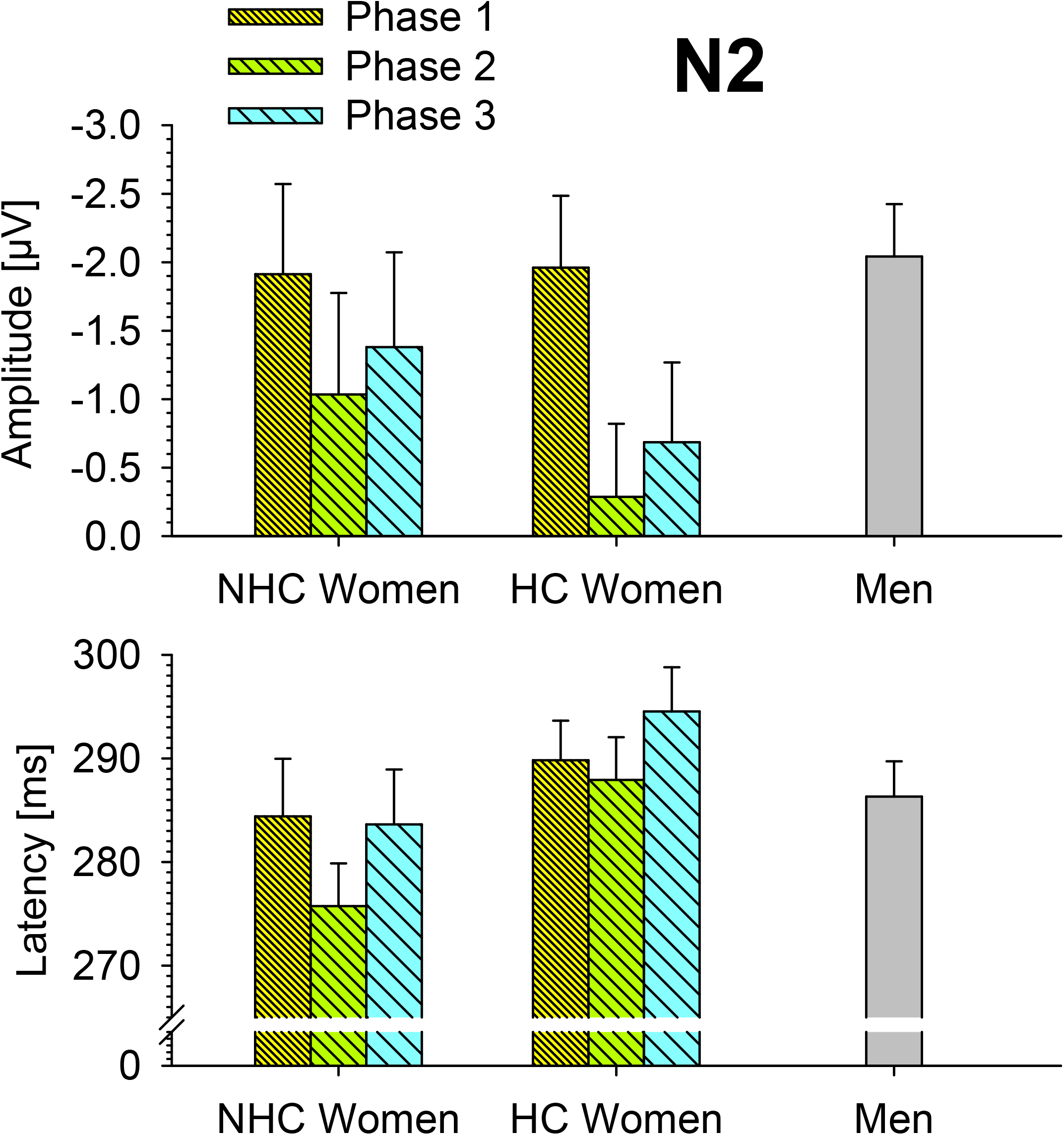
N2 amplitudes (upper panel) and latencies (lower panel) across participant groups and phases at electrode FCz. For NHC and HC groups, results are shown for phases 1, 2, and 3; men were tested once. Error bars, standard errors.

For N2 latency, results of the two-factor ANOVA for NHC and HC groups at the three phases failed to reveal significant effects. There was merely a numerical trend toward slightly longer N2 latencies in HC, than NHC, participants (*F* ≤ 3.66, *p* ≥ 0.06). Comparisons of men and women did not reveal significant differences at any phase (*t* ≤ 0.68, *p* ≥ 0.50).

#### 3.2.2 Electrical imaging

The cortical source of the effect of phase on the N2 amplitude, as found by ERP analysis in women (cf. 3.2.4), was localized using sLORETA. We computed the contrast of phase 1 vs. phase 2 for all women, since the strongest difference in N2 amplitude was obtained for this comparison. These results revealed a focal peak location at MNI coordinates *X* = 35 mm, *Y* = 50 mm, *Z* = 25 mm (*t*_max_ = 4.16, *p* = 0.009, two-tailed) in the region of right superior frontal gyrus (SFS; Brodmann areas, BAs 9/10), with stronger electrical activity in phase 1 compared to phase 2 (Fig. 4). All other comparisons between phases did not reveal significant results.

**Fig. 4.**
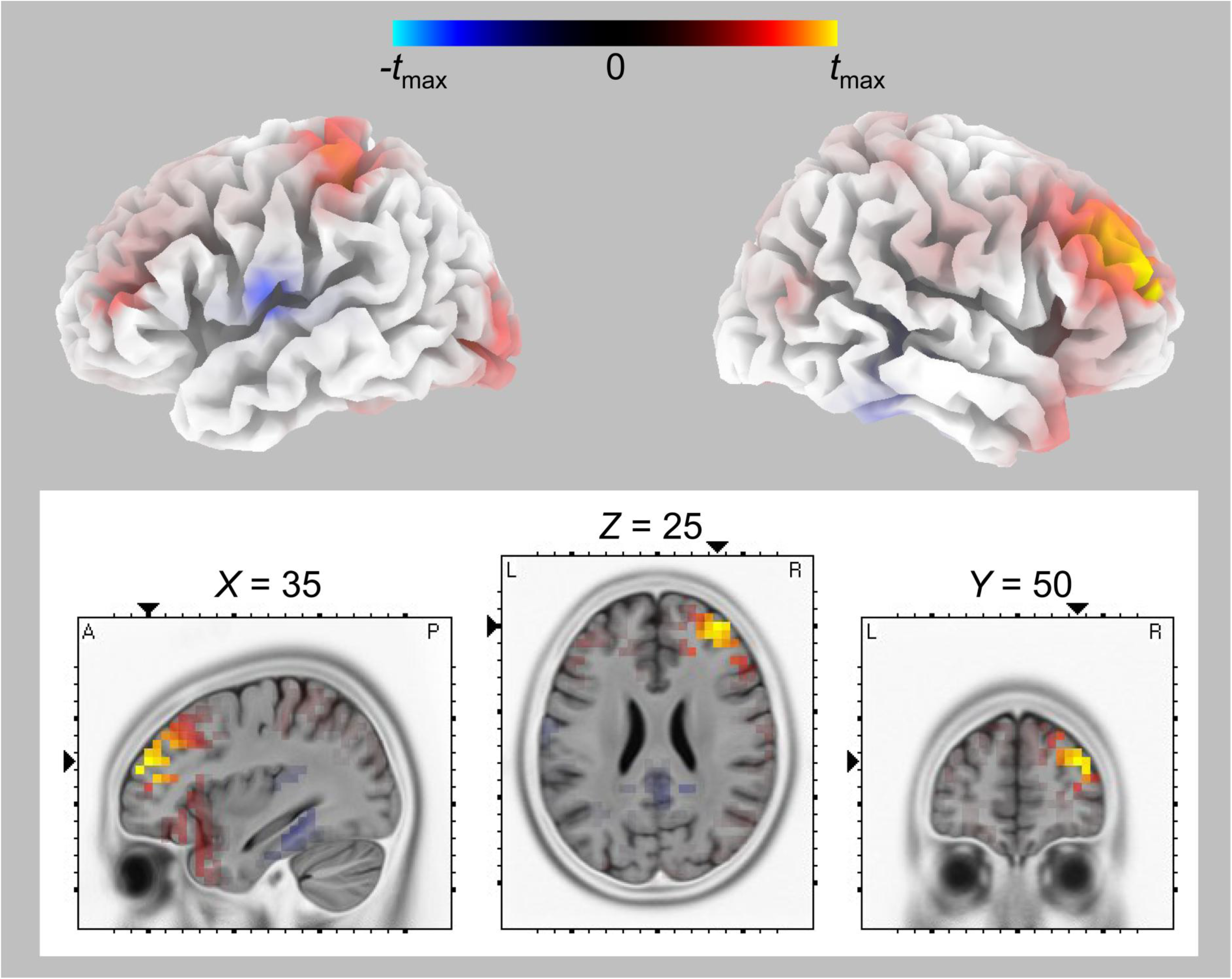
Cortical sources of phase-related differences in women at the time of the N2. Warm colors indicate stronger electrical activity in phase 1 compared to phase 2. The peak activation was located in the region of right superior and middle frontal gyri (MNI coordinates *X* = 35 mm, *Y* = 50 mm, *Z* = 25 mm; *t*_max_ = 4.16, *p* = 0.009, two-tailed). Data are projected onto a single anatomical image with either 3D cortical surface (MRI-template Colin of sLORETA) or sagittal, horizontal, and coronal slices (MRI-template MNI 152-2009c T2 of sLORETA).

In a further analysis, women in phase 2 were compared with men, since a significant difference in N2 amplitude was found for this comparison (cf. 3.2.4). This contrast revealed a peak at MNI coordinates *X* = 55 mm, *Y* = 25 mm, *Z* = 15 mm (*t*_max_ = 3.96, *p* = 0.03, two-tailed) in the region of inferior frontal gyrus (IFG; BA 45), with stronger electrical activity in women than in men (Fig. 5). Comparisons between men and women in other phases did not reveal significant results.

**Fig. 5.**
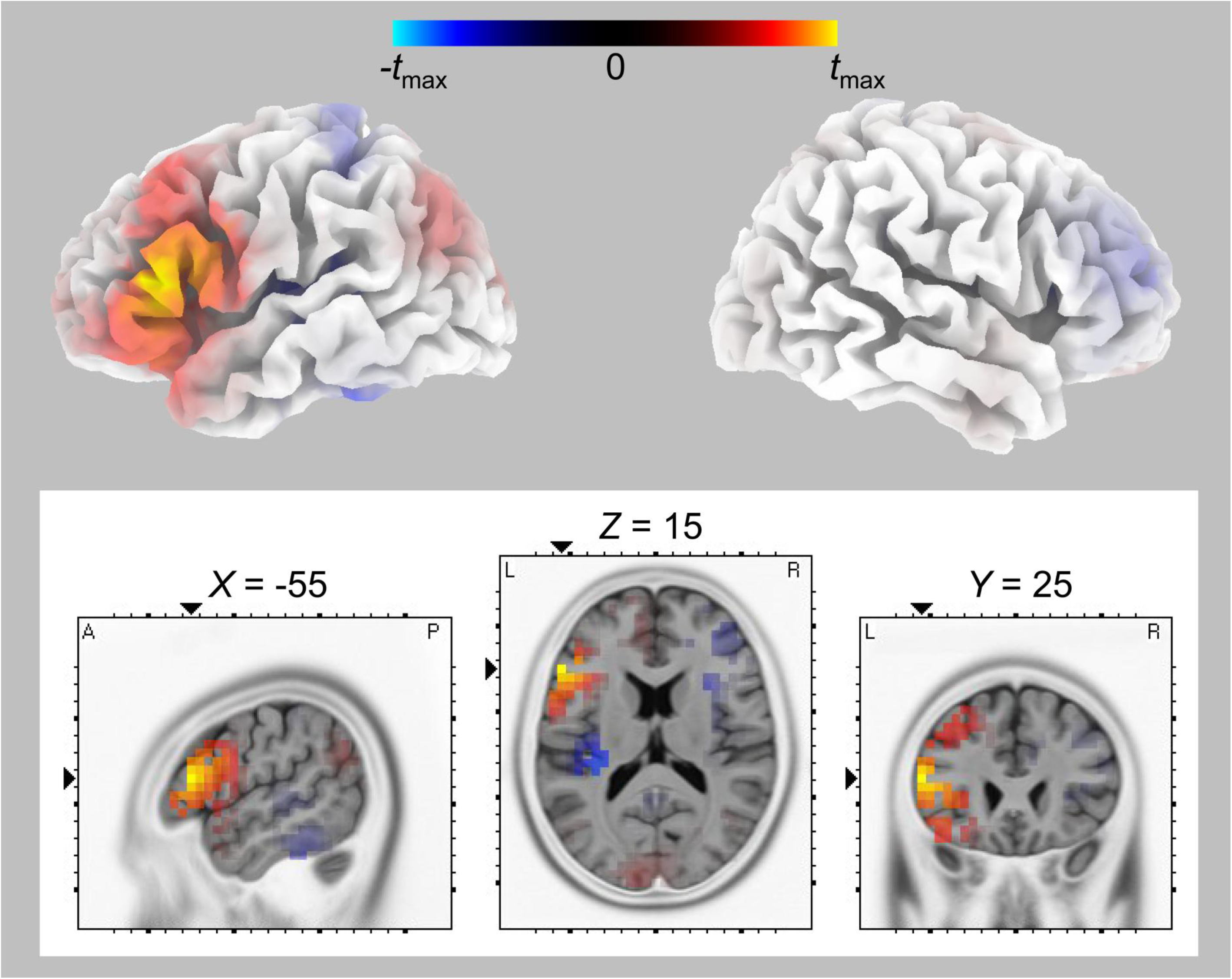
Contrast of women in phase 2 and men at the time of the N2. Source localization analysis revealed a peak in right inferior frontal gyrus (MNI coordinates *X* = 55 mm, *Y* = 25 mm, *Z* = 15 mm; *t*_max_ = 3.96, *p* = 0.03, two-tailed). Warm colors indicate stronger electrical activity in women than in men. Data are projected onto a single anatomical image with either 3D cortical surface (MRI-template Colin of sLORETA) or sagittal, horizontal, and coronal slices (MRI-template MNI 152-2009c T2 of sLORETA).

#### 3.2.3 Secondary ERP analyses

For P1 peak amplitudes at electrode position FCz, the two-factor ANOVA for NHC and HC groups and three phases did not show any significant results (*F* ≤ 0.42, *p* ≥ 0.54; Fig. 6). Also, comparisons of women and men did not indicate significant sex differences in any of the three phases (|*t*| ≤ 0. 71, *p* ≥ 0.48). For P1 latency, the two-factor ANOVA for NHC and HC groups also delivered non-significant results (*F* ≤ 0.44, *p* ≥ 0.61). Moreover, P1 latencies of women and men did not differ in any of the three phases (*t* ≤ 0.20, *p* ≥ 0.84).

**Fig. 6.**
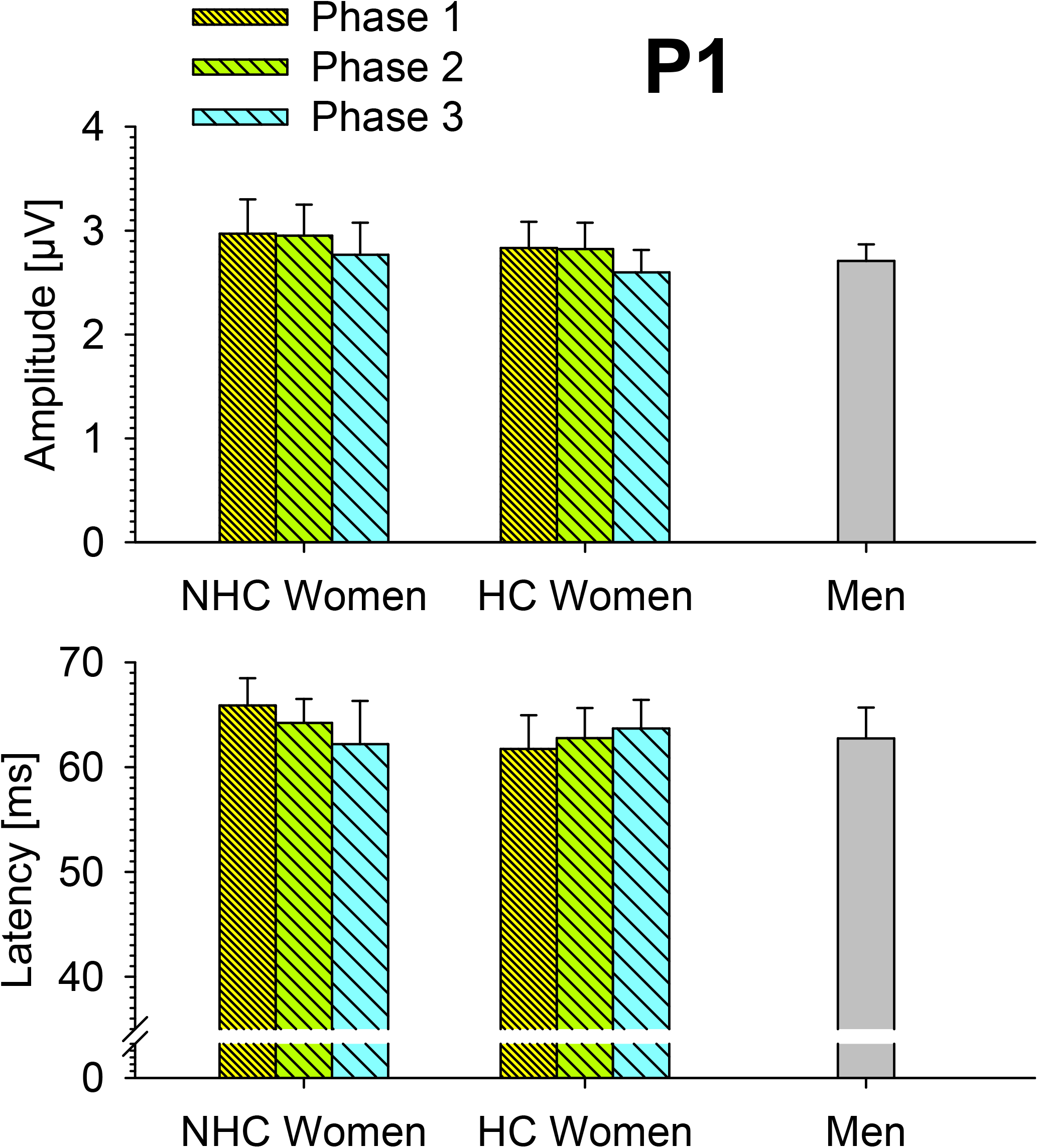
P1 amplitudes (upper panel) and latencies (lower panel) across participant groups and phases at electrode FCz. Conventions are as in Fig. 3.

For N1 peak amplitudes at electrode position Cz, the two-factor ANOVA for NHC and HC groups at three phases did not show any significant results (*F* ≤ 0.29, *p* ≥ 0.75; Fig. 7). Also, no significant sex differences were found at any of the three phases (*t* ≤ 0.99, *p* ≥ 0.33). For N1 latency, results of the two-factor ANOVA for NHC and HC groups also were non-significant (*F* ≤ 1.40, *p* ≥ 0.25). However, the comparisons of women and men indicated significantly longer latencies in men than women in phase 1 (*t*[98] = 3.05, *p* = 0.003, *d* = 0.64) and phase 2 (*t*[98] = 3.46, *p* = 0.0008, *d* = 0.73), but not in phase 3 (*t*[98] = 1.70, *p* = 0.09; Bonferroni-corrected α = 0.017).

**Fig. 7.**
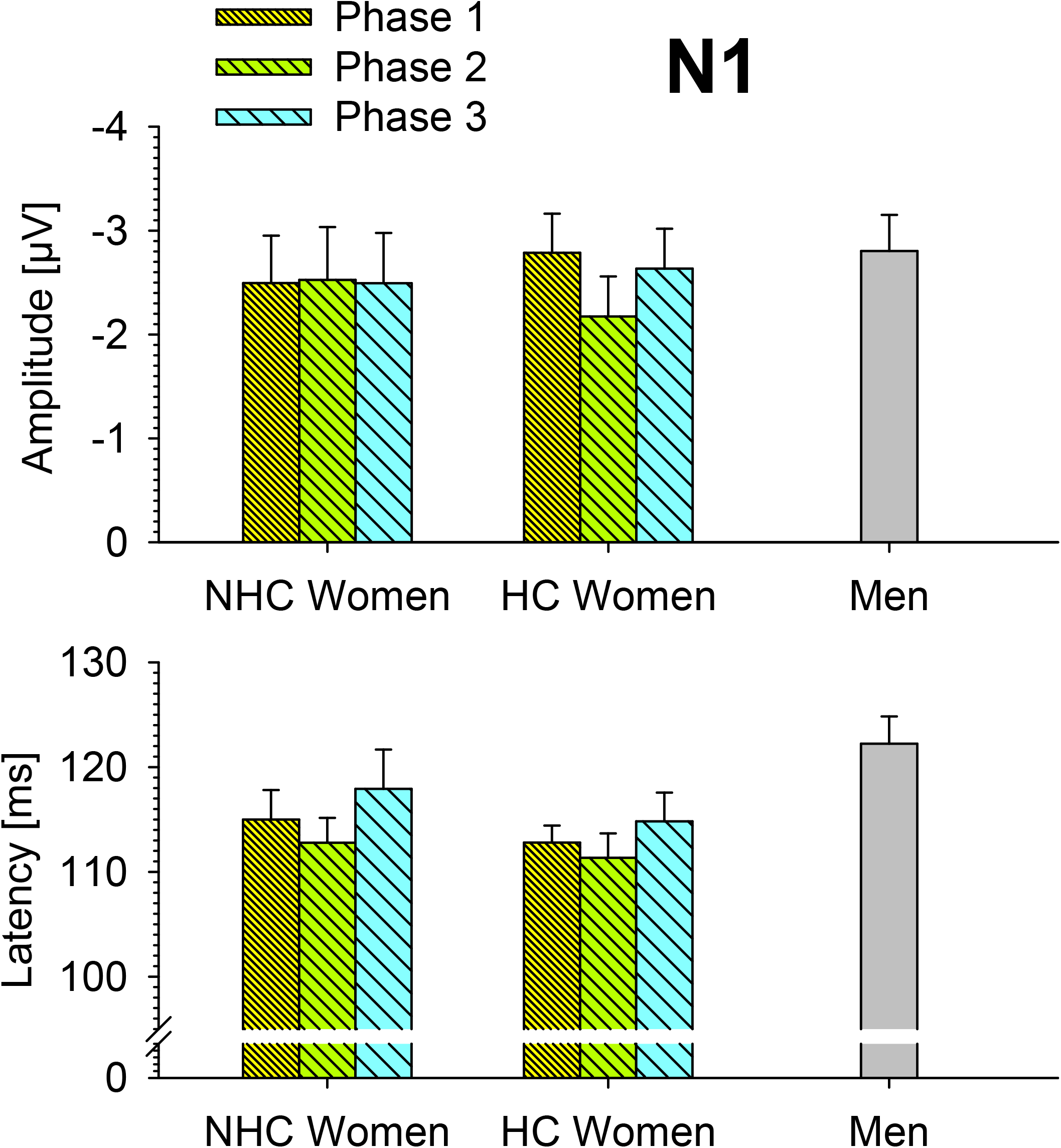
N1 amplitudes (upper panel) and latencies (lower panel) across participant groups and phases at electrode Cz. Conventions are as in Fig. 3.

For P2 peak amplitudes at electrode position Cz, the two-factor ANOVA comparing NHC and HC groups as well as three phases did not reveal significant results (*F* ≤ 1.46, *p* ≥ 0.23; Fig. 8). Comparisons of men and women indicated significantly smaller P2 amplitudes in men than in women in phase 1 (*t*[98] = 2.43, *p* = 0.017, *d* = 0.51), phase 2 (*t*[98] = 3.62, *p* = 0.0005, *d* = 0.76), and phase 3 (*t*[98] = 2.91, *p* = 0.005, *d* = 0.55; Bonferroni-corrected α = 0.017). For P2 latency, results of the two-factor ANOVA for NHC and HC groups were non-significant (*F* ≤ 0.56, *p* ≥ 0.57). Also, comparisons of men and women at the three phases did not show any significant result (|*t*| ≤ 1.82, *p* ≥ 0.07).

**Fig. 8.**
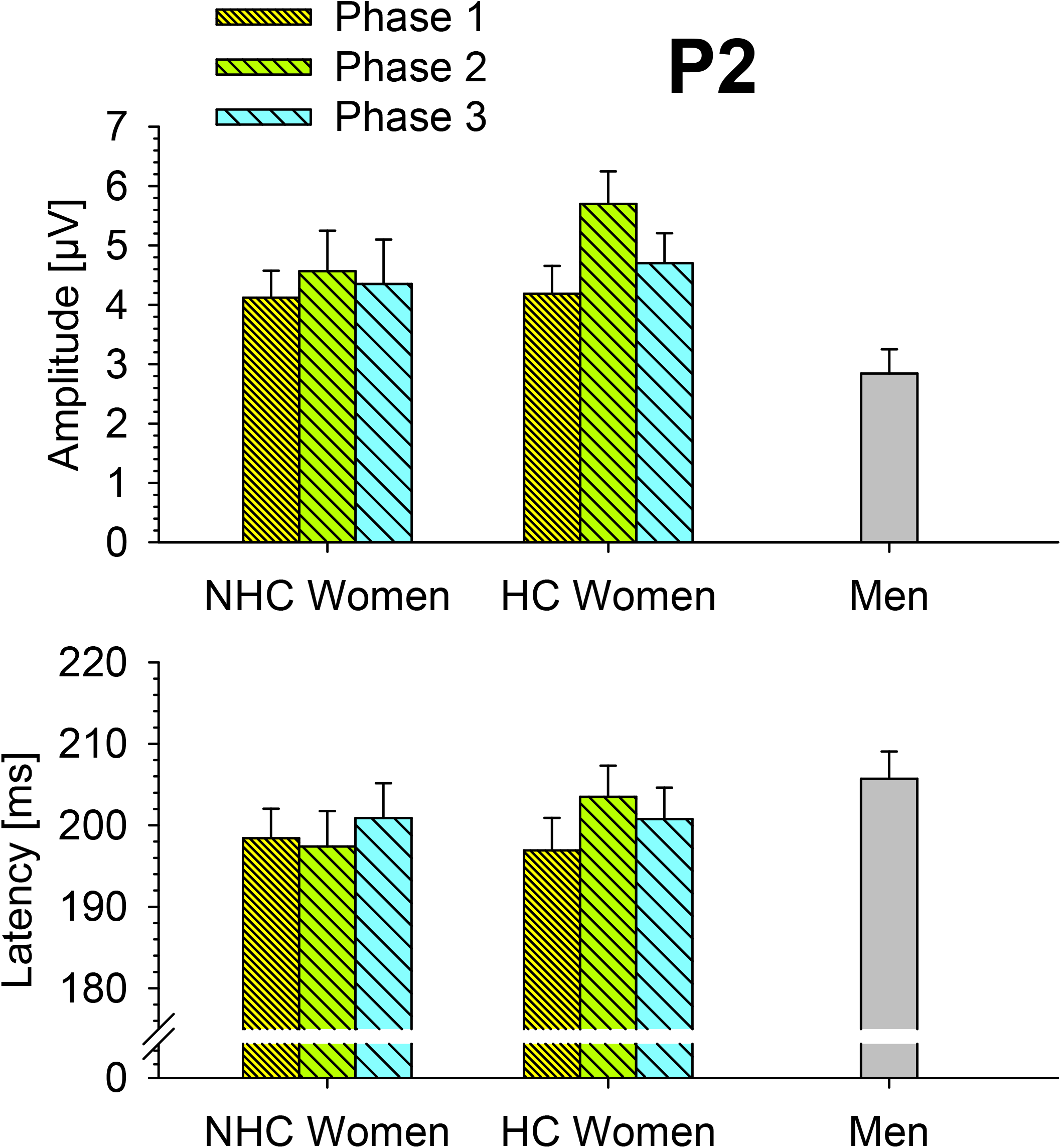
P2 amplitudes (upper panel) and latencies (lower panel) across participant groups and phases at electrode Cz. Conventions are as in Fig. 3.

For LPC peak amplitudes at electrode position Pz, the two-factor ANOVA for NHC and HC groups at three phases did not reveal any significant results (*F* ≤ 1.96, *p* ≥ 0.15; Fig.9). Comparisons of women and men showed a numerical trend of stronger amplitudes in women than men in phase 2, but no significant results (*t* ≤ 1.74, *p* ≥ 0.09). For LPC latency, the two-factor ANOVA did not deliver any significant result (*F* ≤ 0.44, *p* ≥ 0.08). Also, women and men did not significantly differ in any of the three phases, although a numerical trend suggested shorter latencies in women than men in phase 2 (*t* ≤ 1.95, *p* ≥ 0.054).

**Fig. 9.**
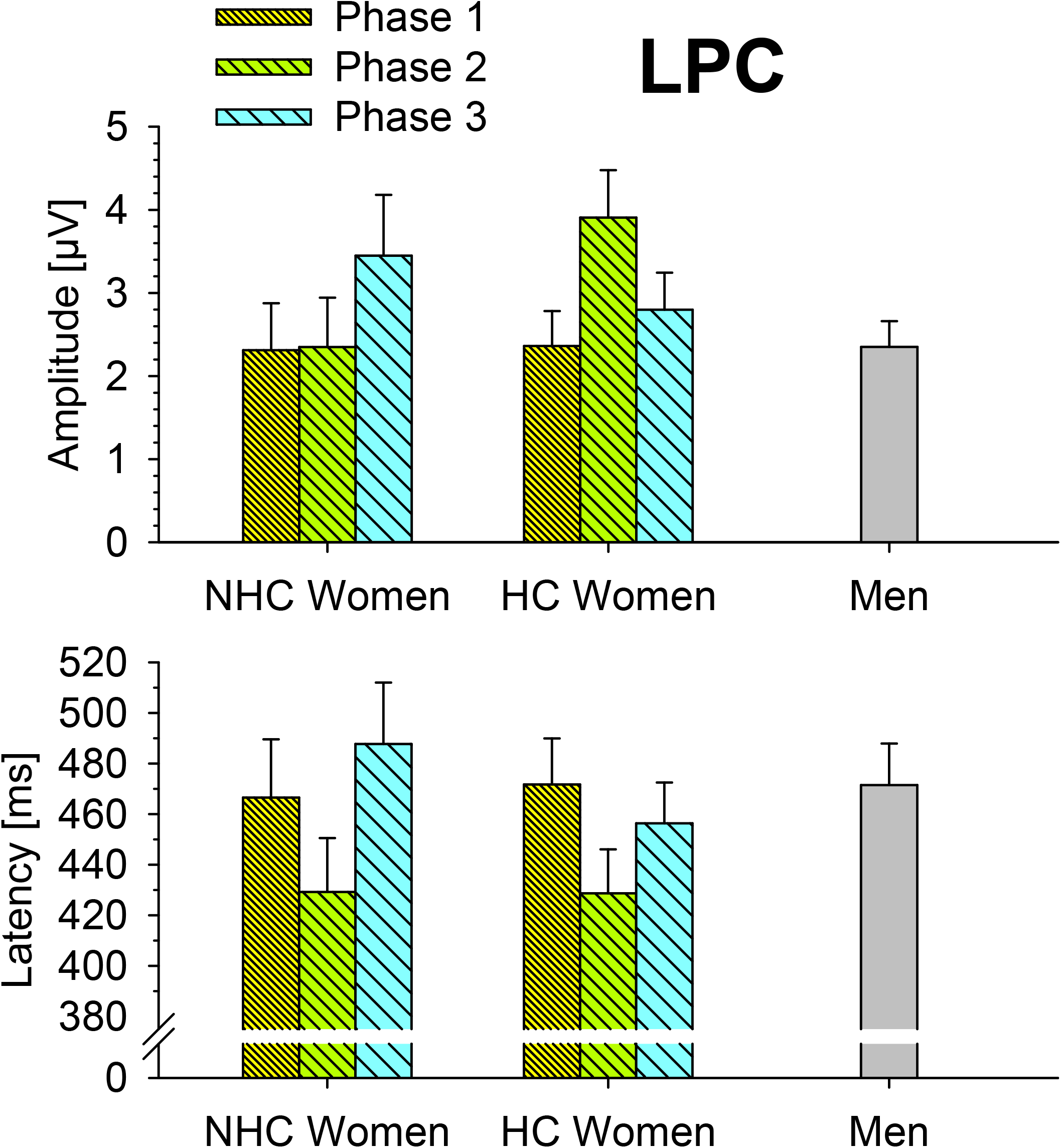
LPC amplitudes (upper panel) and latencies (lower panel) across participant groups and phases at electrode Pz. Conventions are as in Fig. 3.

## 4 Discussion

The present study ought to examine the impact of the hormonal cycle on auditory localization processes. There was a significant variation of the N2 component of the ERP in women depending on the time of recording, with maximum amplitudes in phase 1, minimum amplitudes in phase 2, and intermediate values in phase 3. Comparisons of male and female groups showed significantly lower N2 amplitudes in women than in men during phase 2, but not in other phases. The cyclical variation of the N2 in women was highly specific since all other ERP components analyzed remained largely stable over time. In accordance with these findings, electrical imaging in women at the time of the N2 revealed stronger electrical activity in phase 1 than in phase 2, with the locus of maximum difference in the region of right SFS. On the other hand, the contrast of men versus women in phase 2 indicated slightly, but significantly, lower electrical activity in men than in women in IFG.

The N2 was strongest in the luteal phase of NHC participants (i. e., 7-13 days before the onset of the menstrual cycle) or 7-13 days before the onset of the withdrawal bleeding in HC participants, whereas the numerically weakest N2 amplitudes were found in the phase of menses in NHC participants or between day 1 to 5 of withdrawal bleeding in HC participants. Previous studies on cyclic changes of sex hormone concentrations in the blood of NHC (e.g., Rosenberg & Park, 2002; Walpurger et al., 2004;; Colzato et al., 2010; Tillman, 2010; Sundström-Poromaa, 2018; Zhu et al., 2016; Pletzer et al. 2017; Weis et al., 2019) and HC women (e.g., Gingnell et al., 2013; Montoya & Bos, 2017; Robakis et al., 2019; Menting-Henry et al., 2022) have demonstrated that phase 1, as defined here (i. e., the luteal phase of the natural cycle in NHC women or the initial phase of pill intake in HC women) is characterized by high levels of natural or synthetic estradiol and progesterone, phase 2 (menses or pause of pill intake) by low levels of both these hormones, and phase 3 by high estradiol and low progesterone levels. The observed pattern is therefore compatible with the hormonal differences typically associated with these phases, but the present data cannot determine whether estradiol, progesterone, their combination, or other phase-related factors contributed to the observed N2 modulation.

The statistical analyses did not allow any clear conclusions to be drawn as to which sex hormone plays the main role in this regard. However, the clear maximum in N2 amplitude in phase 1 (cf. Fig. 3) suggests that the combination of high estradiol and progesterone levels was more relevant than a selective effect of estradiol or progesterone alone. In this context, it has to be emphasized that the reduced N2 amplitudes during low hormone phases may not reflect a general impairment but rather a shift towards alternative processing strategies or a different balance between neural systems involved in spatial and non-spatial aspects of auditory perception.

The finding that the effect of phase on the ERP was specific to its N2 component is consistent with the view, based on previous work, that the N2 can be regarded as a correlate of auditory selective spatial attention (e.g., Gamble & Luck, 2011; Gamble & Woldorff, 2014; Lewald & Getzmann, 2015; Lewald et al., 2016; Hanenberg et al., 2019, 2021). That is, sex hormone levels may have specifically modulated brain processes involved in cocktail-party listening. Possibly related modulations of the N2 can also be induced by other factors. Recent studies using similar tasks as in the present approach have also found enhancements of the N2 after short-term spatial audiovisual-congruency training (Hanenberg et al., 2021) and after monopolar anodal transcranial direct-current stimulation (tDCS) of right posterior temporal lobe (Hanenberg et al., 2019). Thus, it is possible that sex hormones target the same processes involved in performing the task as sensory training or tDCS. This view is supported by the results obtained by distributed source analysis. When comparing women across phases, modulation of electrical activity at the time of the N2 was found in right SFS, which was nearby the location of tDCS and training-induced changes in electrical activity reported by Hanenberg et al. (2019, 2021). Given the relatively coarse spatial resolution of the method, this region could also be assigned to the auditory SFS region of the human cortex, as defined by Arnott et al. (2004), which is part of the auditory posterodorsal stream (i. e., posterior superior temporal gyrus, inferior parietal lobule, and SFS) and has been associated with functions of auditory spatial perception and selective attention (Pugh et al., 1996; Nakai et al., 2005; Lee et al., 2012; Braga et al., 2013; Kong et al., 2014; Lewald & Getzmann, 2015; Zündorf et al., 2013, 2014, 2016; Lewald, 2016, 2019; Lewald et al., 2016, 2018). In a more general context, the superior frontal gyrus is known to be involved in higher-order executive processes, including working memory, planning, and complex decision-making (Stuss, 2001; Brass & von Cramon, 2004; Ramnani & Owen, 2004; du Boisgueheneuc et al., 2006). Unlike that, the finding of stronger electrical activity in left IFG in women in phase 2 compared with men seems, at the first glance, unexpected since this region is known to be part of the anteroventral auditory stream, primarily involved in non-spatial functions, such as frequency and pitch processing (e.g., Zatorre et al., 1992; Linden et al., 1999; Alain et al., 2001; Kiehl et al., 2001; Schall et al., 2003; Molholm et al., 2005), auditory working memory (Stevens et al., 2000), and sound identification (for review, see Rauschecker & Tian, 2000; Tranel et al., 2003; Lewis et al., 2004). However, in accordance with the present results, the left IFG has been revealed by fMRI in healthy participants (Zündorf et al., 2013) and voxel-based lesion-behavior mapping in patients with left brain damage (Zündorf et al., 2014) using a similar cocktail-party task as used here. Several further studies have argued that the IFG may be part of a shared non-spatial and spatial cortical auditory network that is involved in both the identification and the localization of sound (Zatorre et al., 2002; Cohen et al., 2005; Lewald et al., 2008; Hill & Miller, 2010; Lewald and Getzmann, 2011; Zündorf et al. 2014, 2016). Also, it is important to note that target localization in the cocktail-party task necessarily required the identification of the target sound among distracters, which might engage both non-spatial and spatial networks. Thus, the present finding of stronger activity in left IFG in women in phase 2 than in men might suggest that the lack of sex hormones in this phase not only reduced dorsofrontal spatial processing in women, but rather resulted in an asymmetrical change in the relationship between activities in left and right auditory cortical streams. That is, the decrease in right dorsofrontal processing in the low-hormone phase 2 was obviously accompanied by stronger engagement of left inferior-frontal processing. Conversely, the elevated hormone levels in phase 1 may have induced a shift toward more intensive processing in the right hemisphere. Since the left IFG was only revealed in contrasts of women vs. men, but not in contrasts of different phases in women, this issue needs, on the one hand, to be examined in further studies. On the other hand, one might speculate that this result could be related to previous findings on variations in lateralization of cognitive functions across the menstrual cycle. Several studies suggested that high levels of estradiol and/or progesterone in follicular and luteal phases reduce the degree of lateralization, as compared to the low-hormone menstrual phase (e.g., Hampson, 1990a; Hampson, 1990b; Sanders and Wenmoth, 1998; Hausmann, 2005; Maki et al., 2002; Hausmann et al., 2002; Hausmann & Güntürkün, 2000; Cowell et al., 2011; Hjelmervik et al., 2012a). In particular, the left-hemispheric language lateralization with right-ear advantage, which typically occurs in dichotic consonant-vowel listening tasks, has been shown to be reduced in the follicular and luteal phases and more pronounced in the menstrual phase (e.g., Hampson, 1990a; Hampson. 1990b; Sanders & Wenmoth, 1998; Hausmann & Güntürkün, 2000; Wadnerkar, Whiteside, & Cowell, 2008; Cowell et al., 2011; Hjelmervik et al., 2012a; Hodgetts, Weis & Hausmann, 2015). Using a musical chord recognition task, Sanders and Wenmoth (1998) found that the typical left-ear advantage for music, indicating right-hemispheric lateralization, was stronger during menses than during the luteal phase. This seems to be in partial opposition to the present results that suggested a more complex pattern of modulation, with a decrease in lateralization of right-hemispheric functions and an apparent increase in lateralization of left-hemispheric functions (that is, a general shift of activity from the right to the left hemisphere) during the low-hormone phase 2 compared with high-hormone phases. However, as already mentioned above, the cocktail-party task used here involved non-spatial (target identification) as well as spatial (target localization) aspects of processing, which could potentially be influenced differently by sex hormones. Comparable studies investigating the effects of the menstrual cycle or sex hormones on neural correlates of auditory attention have so far only been conducted in the non-spatial domain. Using an auditory oddball task (frequency differences of tones), Walpurger et al. (2004) observed significantly longer N2 latencies during the follicular and luteal phases compared to menses, but no modulation of N2 amplitude. In contrast to that, here we found only a non-significant trend for the N2 latency (cf. Fig. 3), but significantly increased N2 amplitudes at related phases. Using an auditory habituation paradigm, Walpurger et al. (2004) also revealed a significant reduction in vertex peak-to-peak potentials during the first block of habituation in the luteal phase, which was correlated with both estradiol and progesterone levels. These authors concluded that hormonal fluctuations across the menstrual cycle may affect earlier ERP components reflecting the cortical arousal response as well as later stages of auditory attentional processing. Although the main result of the present study indicated a specific effect of phase on the N2 amplitude, N1 latencies were also found to be significantly longer in men than women in phases 1 and 2, but not in phase 3. This latter finding might possibly be related to the modulation of N1 amplitude observed in the non-spatial task of Walpurger et al. (2004). Thus, the effect of phase on the N2 amplitude found here could be specific to the spatial domain of auditory attentional processes, as was relevant in the task used.

The present findings in the auditory domain are broadly consistent with previous studies reporting cycle-related variations in visual cognition (for review, see Hausmann et al., 2017) and multisensory perception (Maccora et al., 2023). Significant changes in visual functions depending on menstrual phases have been shown, e. g., for target detection, spatial attention and orientation, and visual sensitivity (for review, see Avitabile et al., 2007; Figueiredo da Mascena et al., 2021; Yao et al., 2022). When MRT was used to investigate cycle-related effects, women showed less activation during tasks involving mental rotation during the mid-luteal phase and increased scores during the menstrual phase, sometimes matching or even surpassing the performance of men (Vandenberg et al., 1978; Voyer et al., 1995; Lippa et al., 2010; Maeda et al., 2013), thus suggesting that low estrogen had a positive impact on visuospatial abilities (Hausmann et al., 2000). At the first glance, our results, which suggest enhanced audiospatial attentional processing at high estrogen levels, seem to be in opposition to these findings from the visuospatial domain. However, quite different mechanisms of neural processing might be involved in both tasks. These results have been primarily associated with the current model that estradiol amplifies N-methyl-D-aspartate (NMDA) glutamate receptor activity while suppressing γ-aminobutyric acid (GABA_A_) transmission, and progesterone enhances inhibition via GABA_A_ receptors (Foy et al., 1999; Hsu & Smith, 2003; Smith, 1994; Smith & Woolley, 2004). High estradiol may thus result in more effective excitatory transmission in cortex. In addition, it has been suggested that the GABAergic system may be critically involved in visual selective attention, with increased GABA_A_ receptor activity widening the attentional focus (Faßbender et al., 2023).

Although the observed pattern of N2 amplitudes is thus compatible with the hormonal profiles typically associated with the phases of the menstrual cycle and the use of hormonal contraceptives, the underlying mechanisms remain unknown. As hormone concentrations and neurotransmitter activity were not measured directly, the present findings do not allow conclusions about the specific contributions of estradiol, progesterone, or their interactions with neurotransmitter systems. This issue might be particularly relevant in the auditory domain, where the effects of sex hormones on neural processing have rarely been studied and are largely unclear (Yadav et al., 2002; Al Mana et al., 2008). As already mentioned above, the findings of stronger electrical activity *(1)* in the right SFS region of women during phase 1 compared with phase 2 and *(2)* in in the left IFG of women in phase 1 compared with men (see above), could be interpreted in the light of previous work on hormonal modulations of functional cerebral asymmetries (e. g., Hausmann & Güntürkün, 2000; Hausmann et al., 2002; Hjelmervik et al., 2012b; for review see Hausmann, 2017; Hidalgo-Lopez et al., 2021). Within the framework of the hormone-receptor interaction model, these studies proposed that excitatory callosal fibers elicit GABA-mediated inhibition in homotopic contralateral regions, with elevated progesterone reducing this interhemispheric inhibition and thereby increasing activation in the nondominant hemisphere. Although estradiol exerts predominantly excitatory effects on glutamatergic receptors – predicting increased interhemispheric inhibition and larger functional cerebral asymmetries – empirical findings indicated enhanced bilateral neural activity at high estradiol levels (Dietrich et al., 2001; Hausmann et al., 2002). Unlike progesterone, estradiol does not acutely affect GABAergic mechanisms (Taubøll et al., 2015), suggesting that reduced interhemispheric inhibition depends on combined action of progesterone on glutamatergic and GABAergic systems, whereas estradiol primarily promotes bilateral excitation. Notably, estrogenic effects on excitability may be context-dependent and may also be inhibitory (Taubøll et al., 2015). On the one hand, the stronger engagement of right SFS during high-hormone phases observed in the present study in women is consistent with the well-established right-hemispheric dominance of spatial attention networks and auditory spatial processing (see above; e. g., Zatorre et al., 1999; Arnott et al., 2004; Spierer et al., 2009; Shulman et al., 2010; Dietz et al., 2014; Zündorf et al., 2014, Hidalgo-Lopez et al., 2021). On the other hand, given that both estradiol and progesterone levels were highly correlated across phases, the present data do not allow for a clear distinction between the respective contributions of progesterone and estradiol to the observed patterns of neural activation. A limitation of the present study is that menstrual cycle phase was determined based on participants’ menstrual history rather than direct measurements of estradiol and progesterone. Although interindividual variability in endogenous hormone concentrations within the assigned cycle phases cannot be excluded, the use of self-reported menstrual history remains, however, a widely accepted approach in cognitive neuroscience research (Allen et al., 2016; Dubol et al., 2021).

From a methodological perspective, these findings provide further support for the emerging consensus that hormonal status must be taken into account in cognitive research on sex differences. The commonly observed advantage of men in visuospatial tasks – a finding from several earlier studies that did not account for the phase of the menstrual cycle among female participants – has increasingly been called into question by more elaborate experiments, which have shown that sex differences due to fluctuations in hormone levels exist only during certain phases of the cycle (Kimura, 1996; Hampson, 1990a; Postma et al., 1999; Hausmann & Güntürkün, 2000; Maki et al., 2002; Hausmann, 2005; Kasai 2021). As suggested by the phase-dependent modulation of the ERP in women found here, this may also apply to sex differences in complex spatial listening tasks, as have been described in previous studies (e. g., Zündorf et al., 2011). Since the paradigm of this study was optimized to investigate changes of the N2 amplitude with correct trials (minimizing contamination of the results by errors), differences in behavioral performance between sexes were absent as expected, while the electrophysiological results revealed significant sex differences depending on phases. In particular, a significant sex difference in N2 amplitude was found exclusively during the menstrual/withdrawal phase. The present results thus support the view that sex-related differences in auditory attentional brain processing are more likely attributable to fluctuating hormonal factors rather than to a constant advantage for men.

## Acknowledgements

This work was supported by the Wilhelm and Günter Esser Foundation. The authors wish to thank Dagmar Bieneck, Florian Reiche, and Jonathan Schuchert for help in running the experiments, and Peter Dillmann for preparing software and parts of the electronic equipment.

## Notes

### Competing Interest Statement

The authors have declared no competing interest.

### Summary of Updates

abstract and text updated to clarify, order of the figures changed

